# Screening of hydrocarbon-stapled peptides for inhibition of calcium-triggered exocytosis

**DOI:** 10.1101/2022.03.21.484632

**Authors:** Ying Lai, Michael J. Tuvim, Jeremy Leitz, John Peters, Richard A. Pfuetzner, Luis Esquivies, Qiangjun Zhou, Barbara Czako, Jason B. Cross, Philip Jones, Burton F. Dickey, Axel T. Brunger

## Abstract

The so-called primary interface between the SNARE complex and synaptotagmin-1 (Syt1) is essential for Ca^2+^-triggered neurotransmitter release in neuronal synapses. The interacting residues of the primary interface are conserved across different species for synaptotagmins (Syt1, Syt2, Syt9), SNAP-25, and syntaxin-1A homologs involved in fast synchronous release. This Ca^2+^-independent interface forms prior to Ca^2+^-triggering and plays a role in synaptic vesicle priming. This primary interface is also conserved in the fusion machinery that is responsible for mucin granule membrane fusion. Ca^2+^-stimulated mucin secretion is mediated by the SNAREs syntaxin-3, SNAP-23, VAMP8, synaptotagmin-2, and other proteins. Here, we designed and screened a series of hydrocarbon-stapled peptides consisting of SNAP-25 fragments that included some of the key residues involved in the primary interface as observed in high-resolution crystal structures. We selected a subset of four stapled peptides that were highly α-helical as assessed by circular dichroism and that inhibited both Ca^2+^-independent and Ca^2+^-triggered ensemble lipid-mixing with neuronal SNAREs and Syt1. In a single-vesicle content-mixing assay with reconstituted neuronal SNAREs and synaptotagmin-1 or with reconstituted airway SNAREs and synaptotagmin-2, the selected peptides also suppressed Ca^2+^-triggered fusion. Taken together, hydrocarbon-stapled peptides that interfere with the primary interface consequently inhibit Ca^2+^-triggered exocytosis. Our inhibitor screen suggests that these compounds may be useful to combat mucus hypersecretion that is a major cause of airway obstruction in the pathophysiology of COPD, asthma and cystic fibrosis.

## Introduction

SNARE-mediated membrane fusion is a ubiquitous mechanism for many biological processes, wherein zippering of the SNARE complex drives membranes together and leads to membrane fusion (Brunger et al., 2018; Rothman, 2014; Südhof, 2013). For Ca^2+^-triggered membrane fusion, SNAREs must cooperate with Ca^2+^ sensors. For example, the neuronal SNARE complex (consisting of synaptobrevin-2/VAMP-2, syntaxin-1A (Stx1), and SNAP-25A) forms a specific interface with the Ca^2+^ binding C2B domain of synaptotagmin-1 (Syt1) (Zhou et al., 2017, 2015). The primary interface is conserved in fast synchronous neurotransmitter release (Zhou et al., 2017, 2015). This interface forms prior to Ca^2+^ triggering during synaptic vesicle priming. Ca^2+^ binding to Syt1 remodels or dissociates the primary complex, likely also involving membrane deformation to promote fusion (Zhou et al., 2017, 2015). Disruption of the primary SNARE– Syt1 complex interface by mutations abolished fast synchronous release in cultured neurons and greatly reduced the Ca^2+^-triggered amplitude in single-vesicle fusion experiments. Moreover, the primary interface has been observed in solution by nuclear magnetic resonance (NMR) and fluorescence spectroscopy experiments (Voleti et al., 2020), and it is stabilized by interactions with the plasma membrane.

The primary interface is predicted to be conserved well beyond the neuronal exocytosis machinery (Zhou et al., 2015). Here, we additionally focus on the airway epithelial exocytic machinery because of the role of Ca^2+^-stimulated secretion in lung pathobiology. Rapid secretion of highly produced mucins overwhelms the hydration capacity of the airways, resulting in concentrated mucus that is excessively viscoelastic and adhesive. Such concentrated mucus cannot be propelled by ciliary beating, adheres to the airway wall, and occludes the airway lumen. This results in airflow obstruction and lung inflammation in common lung diseases including asthma, chronic obstructive pulmonary disease (COPD), and cystic fibrosis (Boucher, 2019; Fahy and Dickey, 2010). Airway mucins are secreted both at a low baseline rate that promotes effective mucociliary clearance, and a high stimulated rate that can lead to the pathology described above. In recent years, molecular components mediating these two modes of secretion have been identified (Davis and Dickey, 2008; Fahy and Dickey, 2010; Jaramillo et al., 2018) and as in neurons, these include SNAREs, and Munc18, Munc13, and synaptotagmins, as follows.

Both baseline and stimulated mucin secretion are impaired in SNAP-23 heterozygous mutant mice (Ren et al., 2015), VAMP8 knock-out mice (Jones et al., 2012), and Munc13-2 knock-out mice (Zhu et al., 2008). In contrast, stimulated mucin secretion is selectively impaired in Munc18-2 and Syt2 deletant mice (Jaramillo et al., 2019; Kim et al., 2012; Tuvim et al., 2009), whereas baseline secretion is selectively impaired in Munc18-1 deletant mice (Jaramillo et al., 2019). Syntaxin-3 (Stx3) binds and colocalizes with Munc18-2 in barrier epithelia (Riento et al., 1998), and immunoprecipitation of Stx3 in airway secretory cells pulls down Munc18-2 (Kim et al., 2012). Thus, we hypothesize that the primary interface in stimulated mucin secretion forms between synaptotagmin-2 (Syt2) and the airway SNARE complex consisting of VAMP8, Stx3, and SNAP-23. Antagonism of this interaction while leaving homeostatic baseline mucin secretion intact could have therapeutic value in muco-obstructive lung diseases.

Many of the residues that are involved in the primary neuronal interface are contributed by the N-terminal α-helix of SNAP-25 that forms one of the four α-helices in ternary the SNARE complex (Sutton et al., 1998; Zhou et al., 2015). Theoretically, peptides derived from this region of SNAP-25 could be used to specifically interfere with the primary SNARE–Syt1 interface. However, small peptides are typically unstructured and their efficacy as an inhibitor may be inefficient in the absence of helical structure. In this work, to induce α-helical structure, we use non-natural amino acids containing olefin-bearing groups to generate hydrocarbon-stapled peptides by a ring-closing metathesis reaction using a Grubbs catalyst. We designed 12 stapled peptides and tested them with lipid-mixing fusion assays. A subset of these stapled peptides that exhibited the highest α-helicity disrupted Ca^2+^-triggered exocytosis using single-vesicle content-mixing assays with either reconstituted neuronal SNAREs and Syt1 or with reconstituted airway SNAREs and Syt2. Taken together, our hydrocarbon-stapled peptides can be used to reduce both stimulated neurotransmitter release and mucin secretion.

## Results

### Design and characterization of stapled peptides

Since the primary interface is critical for Ca^2+^-triggered exocytosis (Zhou et al., 2015), it should be possible to interfere with this process by disrupting this protein-protein interface with a suitable compound. We designed a series of hydrocarbon-stapled peptides with varying lengths (16-22 amino acids), named SP1-SP12, that cover various portions of this interface of the N-terminal α-helix of SNAP-25A that is involved in the primary interface with Syt1 (Figs. 1A, 1B). The positions of non-natural amino acid substitutions *(S5* and *R8* olefinic sidechains) were chosen to be away from the primary interface (Fig. 1C). We also used slightly different start and end positions within the SNAP-25A sequence that interacts with Syt1 (Fig. 1B). The hydrocarbon-staples were made to bridge three *(S5* substitution positions *i* and *i*+4) or six *(R8* substitution positions *i* and *i*+7) amino acids within the peptides. As a control, we used a linear peptide (P0) that corresponds to residues 37-58 of SNAP-25A. Liquid chromatography mass spectrometry of the synthesized peptides indicated molecular weights that were close to their predicted molecular weights (Supplementary Figs. S1-S12), suggesting that the synthesized peptides have the correct sequence and properly formed staples.

**Figure 1.**
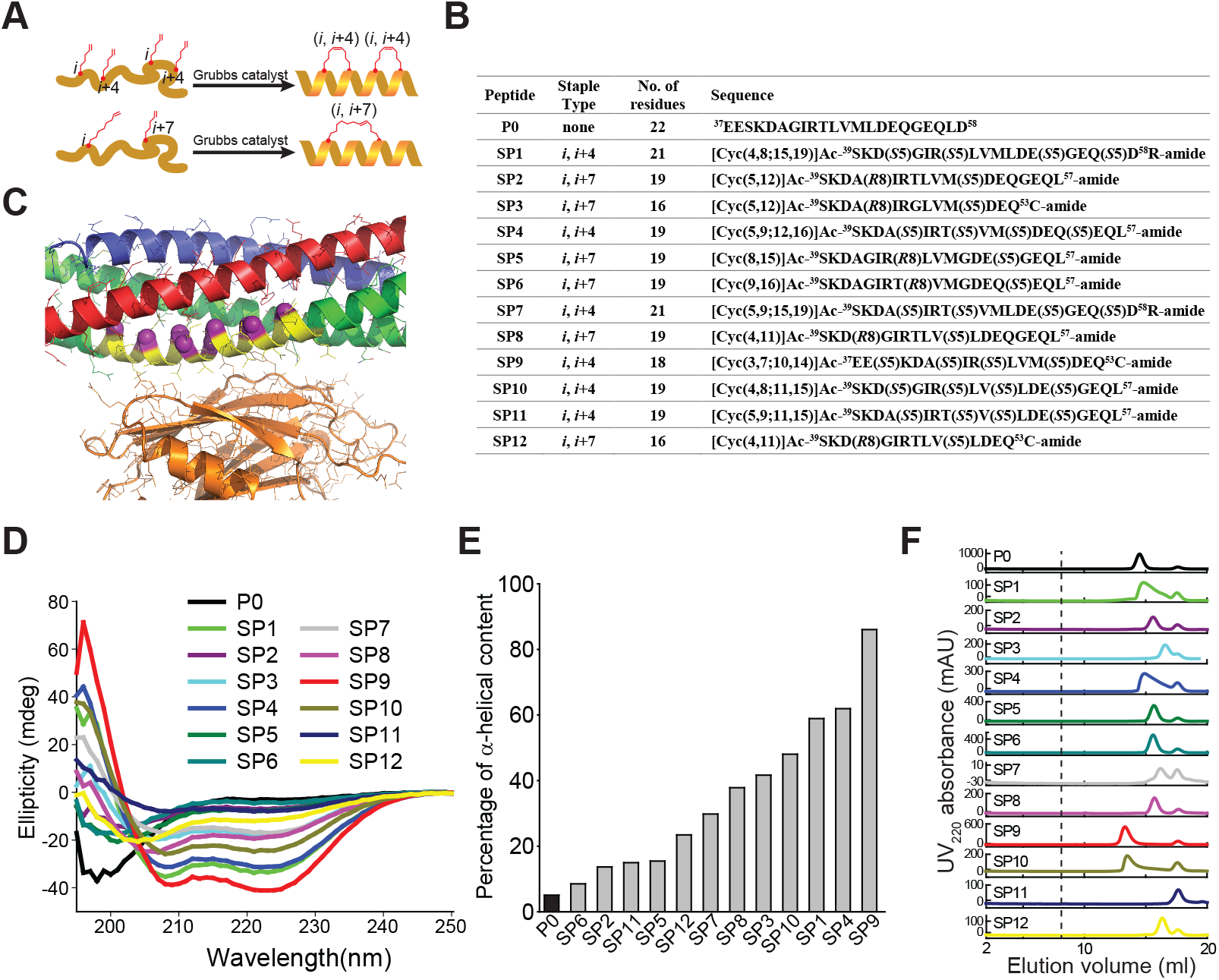
Design and characterization of stapled peptides. **A,** Schematic diagram for stapled peptides synthesis. Hydrocarbon-stapled peptides are formed via cross-linking the residues at the specified positions. P0 is the wildtype sequence for SNAP-25A, residues 37-58. **B,** Design of synthesized stapled peptides with varying lengths of olefinic side chains and different positions of the non-natural amino acid substitutions. The superscripts specify the starting and end positions within the SNAP-25A sequence. *Sn* or *Rn* indicates *S* or *R* stereochemistry at the α-carbon, respectively, *n* indicates the number of carbon atoms in the olefinic side chains. **C**, Close-up view of the primary interface between the neuronal SNARE complex (VAMP-2-blue, Stx1 - red, SNAP-25A - green) and the C2B domain of Syt1 (orange) (PDB ID 5W5C), indicating the region (yellow) from which the peptides were chosen. The Cα-positions of the *S5* and *R7* non-natural amino acid substitutions of all the designs are indicated by purple spheres. **D**, CD spectra of the specified peptides measured at 100 mM concentration, pH 7.4, 25 ± 1 °C. **E**, Percentage of α-helical content of the specified peptides as estimated by dividing the mean residue ellipticity [φ]222_obs_ by the reported [φ]222_obs_ for a model helical decapeptide. **F**, Size exclusion chromatography (SEC) profiles of the specified peptides. Each peptide was filtered with a 0.2 micrometer filter and then loaded on a Superdex 75 column in buffer V (20 mM HEPES, pH 7.4, 90 mM NaCl). The dashed line indicates the border of the void volume at ~8 ml.

Circular dichroism spectroscopy revealed that the non-stapled P0 peptide displays only 5% α-helicity in solution and presumably forms a random coil (Fig. 1D). In contrast, many of the stapled peptides had substantially higher α-helical content. Some of the stapled peptides with two hydrocarbon staples have the highest α-helicity (in particular, SP1, SP4, SP9, and SP10), with an α-helical content up to 86% (Fig. 1E). Note, however, that one of the stapled peptides with two hydrocarbon staples has lower α-helicity (SP11). None of the peptides aggregated as assessed by size exclusion chromatography (Fig. 1F); the differences in retention times are likely related to differences in shape or possible interactions with the resin of the size exclusion column.

**Figure 2.**
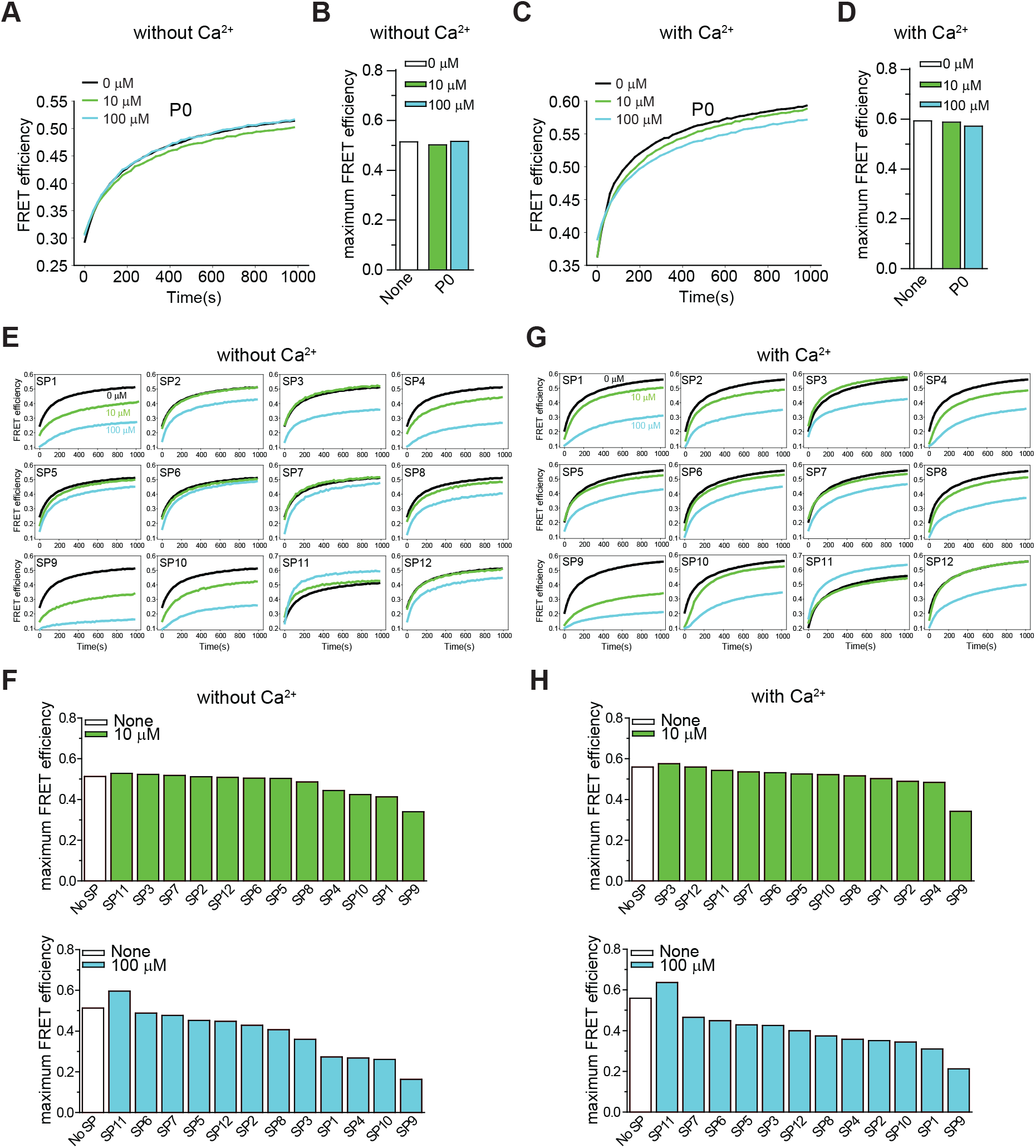
Effect of stapled peptides in ensemble lipid mixing assays with neuronal SNAREs and Syt1. **A**, The non-stapled peptide P0 has little effect on Ca^2+^-independent ensemble lipid mixing measurements of vesicle-vesicle fusion with SV and PM vesicles (Methods). The two groups of vesicles are mixed at the same molar ratio with a final lipid concentration of 0.1 mM. The time traces show FRET efficiency upon mixing dye-labeled neuronal PM- and SV-vesicles (Methods). The black line is a control without peptide. The green and blue lines indicate the lipid mixing experiments with 10 μM and 100 μM P0, respectively. **B**, Corresponding maximum FRET efficiency within the observation period of 1000 sec. **C**, P0 has little effect on Ca^2+^-triggered ensemble lipid mixing measurements of vesicle-vesicle fusion. The two groups of vesicles are mixed at the same molar ratio with a final lipid concentration of 0.1 mM. The time traces show FRET efficiency upon mixing dye-labeled neuronal PM- and SV-vesicles (Methods). The black line is a control without peptide. The green and blue lines represent the reaction with 10 μM and 100 μM P0, respectively. **D,** Corresponding maximum FRET efficiency within the observation period of 1000 sec. **E,** Effect of stapled peptides in absence of Ca^2+^. The two groups of vesicles (SV and PM) were mixed at the same molar ratio with a final lipid concentration of 0.1 mM. The time traces show FRET efficiency upon mixing neuronal dye-labeled PM- and SV-vesicles (Methods). **F,** Corresponding maximum FRET efficiency within the observation period of 1000 sec. **G,** Effect of stapled peptides in the presence of Ca^2+^. The two groups of vesicles (SV and PM) were mixed at the same molar ratio with a final lipid concentration of 0.1 mM. The time traces show FRET efficiency upon mixing neuronal PM- and SV-vesicles. **H,** Corresponding maximum FRET efficiency within the observation period of 1000 sec.

### Screening of stapled peptides with a lipid-mixing assay

We first tested the effects of the peptides with an ensemble lipid mixing (fusion) assay with reconstituted neuronal SNAREs and Syt1. Two types of vesicles were reconstituted: plasma-membrane mimicking vesicles containing 1% DiI with reconstituted Stx1 and SNAP-25A (referred to as PM vesicles) and vesicles containing 1% DiD with reconstituted synaptobrevin-2 and Syt1 that mimic synaptic vesicles (referred to as SV vesicles). The two groups of vesicles were mixed at equal molar ratio with final lipid concentration of 0.1 mM, and fluorescence changes of DiI and DiD were recorded. As expected, addition of the non-stapled peptide P0 has little effect on both Ca^2+^-independent and Ca^2+^-triggered lipid mixing, while many of the stapled peptides reduced lipid mixing both in the absence and presence of Ca^2+^ (Fig. 2). The stapled peptides SP1, SP4, SP9, and SP10 consistently exhibited large inhibition in the lipid mixing assays. Since these four peptides also have the largest α-helical content (Fig. 1D), this suggests that the helical conformation of the peptides is essential for their inhibitory activity.

### Screening of stapled peptides with a single vesicle content-mixing assay

The ensemble lipid mixing assay is relatively fast and allows one to screen many peptides. However, ensemble lipid mixing experiments have considerable caveats. For example, lipid mixing can occur in the absence of full fusion owing to metastable hemifusion diaphragms, and FRET efficiency measurements are influenced by differences in vesicle-vesicle association (docking), potentially obscuring the effects of the peptides. A better surrogate for both neurotransmitter release and mucin secretion is a single-vesicle content-mixing assay since it accurately measures content exchange and it is capable of distinguishing between vesicle association, lipid mixing, and content mixing (Diao et al., 2012; Kyoung et al., 2011; Lai et al., 2022). Since a single-vesicle content mixing assay is considerably more complicated and time-consuming, we only tested the subset of the four stapled peptides (SP1, SP4, SP9, SP10) that consistently showed the largest effects in ensemble lipid mixing experiments (Fig. 3A). In the single-vesicle content-mixing assay, SV vesicles are labeled with a soluble fluorescent content dye (sulforhodamine B) at a moderately self-quenching concentration. Content mixing between SV and PM vesicles results in an increase in volume, and consequently in an increase of content-dye fluorescence intensity. Thus, this assay allows precise measurement of content mixing. Vesicle association was measured by the stable association of a SV vesicle with a surface-tethered PM vesicle, and content mixing data were normalized with respect to the number of associated vesicles. Vesicle association can be influenced by *trans*-SNARE complex formation, Syt1/Syt2–SNARE complex interactions, Syt1/Syt2–syntaxin-SNAP25 subcomplex interactions, and Syt1/Syt2–membrane interactions.

**Figure 3.**
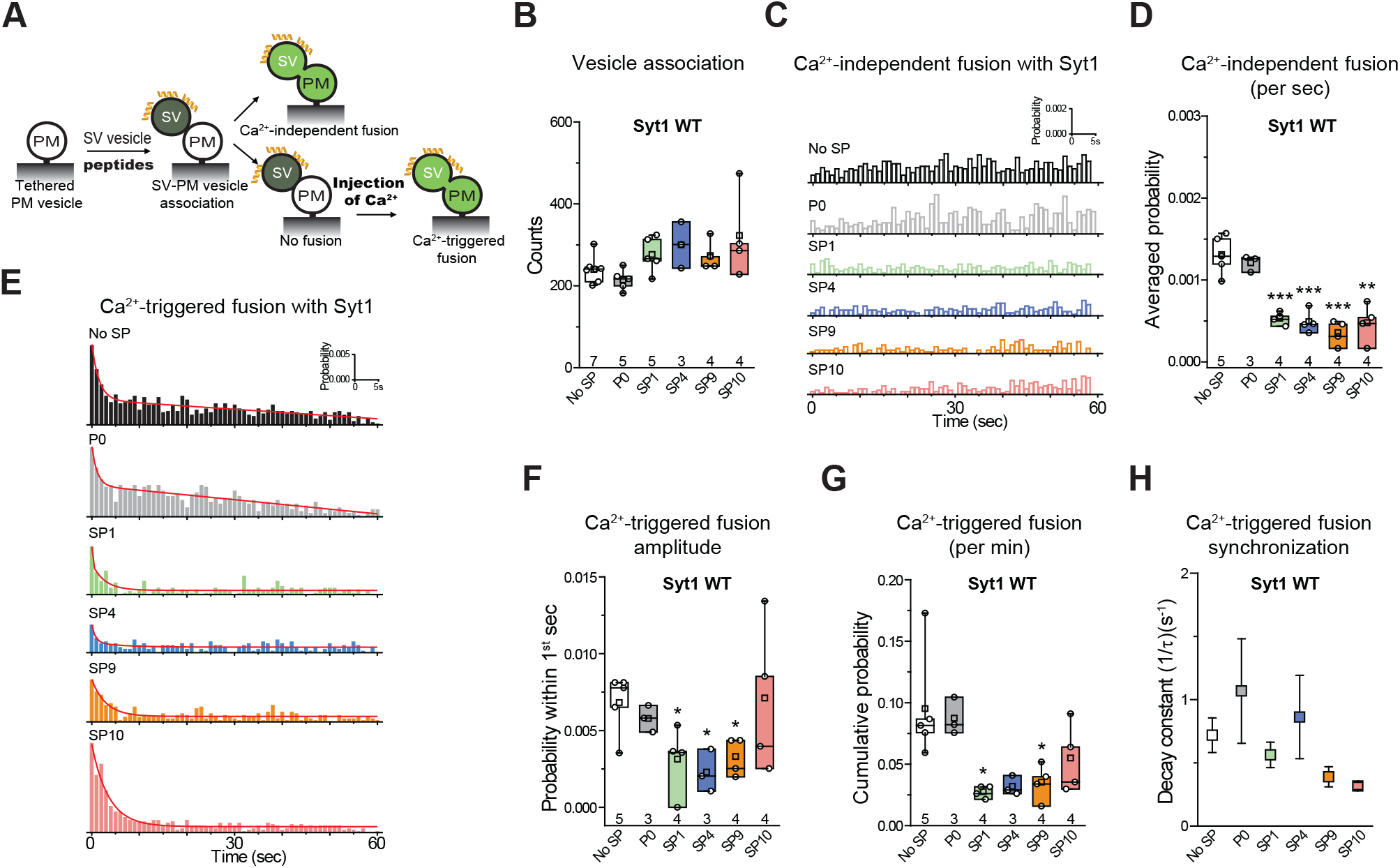
Stapled peptides inhibit both Ca^2+^-independent and Ca^2+^-triggered content mixing with reconstituted neuronal SNAREs and Syt1. **A,** Schematic of the single-vesicle content-mixing assay. Neuronal PM: plasma membrane mimic vesicles with reconstituted Stx1A and SNAP-25A; SV: synaptic vesicle mimic with reconstituted VAMP2 and Syt1. After SV - neuronal PM vesicle association, vesicle pairs either undergo Ca^2+^-independent fusion or remain associated until fusion is triggered by Ca^2+^ addition. 10 μM of each specific stapled peptide was added together with SV vesicles and was present in all subsequent stages. For details of the reconstitution and lipid compositions, see Methods in (Lai et al., 2022). **B,** Effects of stapled peptides on vesicle association. **C,** Corresponding Ca^2+^-independent fusion probabilities. **D,** Corresponding average probabilities of Ca^2+^-independent fusion events per second (left to right: *** p = 0.00044, *** p = 0.00051, *** p = 0.00022, ** p = 0.0012). **E,** Corresponding Ca^2+^-triggered fusion probabilities. (**F-H**) Corresponding Ca^2+^-triggered fusion amplitudes of the first 1-sec time bin upon 500 μM Ca^2+^-injection (**F**) (left to right: * p = 0.034, * p = 0.013, * p = 0.017), the cumulative Ca^2+^-triggered fusion probability within 1 min (**G**) (left to right: * p = 0.021, * p = 0.039), and the decay rate (1/τ) of the Ca^2+^-triggered fusion histogram (**H**). The fusion probabilities and amplitudes were normalized to the number of analyzed neuronal SV - neuronal PM vesicle pairs (Supplementary Table S1). Panels **B**, **D**, **F**, **G** show box plots and data points for n (indicated below each box plot) independent repeat experiments (Supplementary Table S1): the whiskers show the min and max values (excluding outliers), the box limits are the 25th and 75th percentiles, the square point denotes the mean value, and the center line denotes the median. Decay constants (boxes) and error estimates (bars) in panels **H** computed from the covariance matrix upon fitting the corresponding histograms combining all repeats with a single exponential decay function using the Levenberg-Marquardt algorithm. * p < 0.05, ** p < 0.01, *** p < 0.001 by Student’s t-test, compared to the experiment without the stapled peptides.

As expected, inclusion of 10 μM P0 in the fusion assay had no effect on either Ca^2+^-independent or Ca^2+^-triggered fusion (Fig. 3C-H). In contrast, the four stapled peptides reduced both the cumulative Ca^2+^-independent fusion (up to 60%) and Ca^2+^-triggered fusion (up to 70%) probabilities (Figs. 3C-H), but they had little effect on vesicle association (Fig. 3B). When Syt1 was omitted from the reconstituted SV vesicles, none of the four stapled peptides showed any effect on Ca^2+^-independent fusion (Fig. 4A-C), corroborating the notion that the effect of the stapled peptides depends on Syt1. Moreover, there is no effect of the stapled peptides on Ca^2+^-independent fusion with SNAREs only (Figure 4A-C), suggesting that SNARE complex formation is not affected by the presence of the stapled peptides.

**Figure 4.**
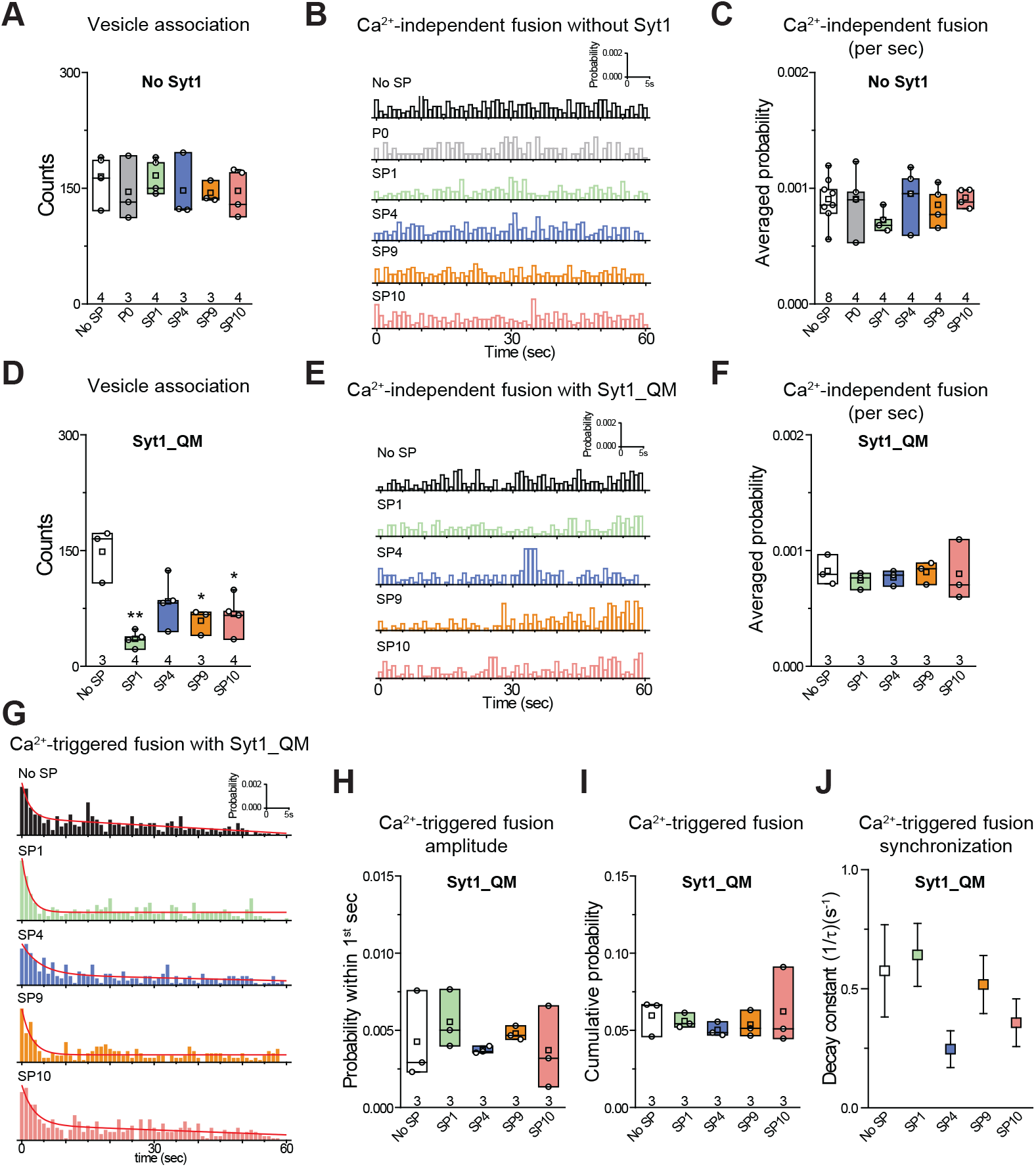
The quintuple Syt1_QM mutant and omission of Syt1 alleviate the inhibitory effects of stapled peptides. **A,** Effects of 10 μM of each of the specified peptides on vesicle association. **B,** Corresponding Ca^2+^-independent fusion probabilities. **C,** Corresponding average probabilities of Ca^2+^-independent fusion events per second. **D,** Effects of 10 μM of each of the specified peptides on vesicle association (left to right: ** p = 0.0016, * p = 0.016, * p = 0.017). **E,** Corresponding Ca^2+^-independent fusion probabilities. **F,** Corresponding average probabilities of Ca^2+^-independent fusion events per second. **G,** Corresponding Ca^2+^-triggered fusion probabilities. (**H-J**) Corresponding Ca^2+^ -triggered fusion amplitude of the first 1-sec time bin upon 500 μM Ca^2+^-injection (**H**), the cumulative Ca^2+^-triggered fusion probability within 1 min (**I**), and the decay rate (1/τ) of the Ca^2+^-triggered fusion histogram (**J**). The fusion probabilities and amplitudes were normalized to the number of analyzed neuronal SV - neuronal PM vesicle pairs (Supplementary Table S1). Panels **A**, **C**, **D**, **F**, **H**, **I** show box plots and data points for n (indicated below each box plot) independent repeat experiments (Supplementary Table S1): the whiskers show the min and max values (excluding outliers), the box limits are the 25th and 75th percentiles, the square point denotes the mean value, and the center line denotes the median. Decay constants (boxes) and error estimates (bars) in panels **J** computed from the covariance matrix upon fitting the corresponding histograms combining all repeats with a single exponential decay function using the Levenberg-Marquardt algorithm. * p < 0.05, ** p < 0.01 by Student’s t-test, compared to the experiment without the stapled peptides.

We next asked the question whether the inhibition of the fusion probability by the stapled peptides is due to specific interference with the interaction between the neuronal SNARE complex and Syt1 via the primary interface. We performed the single-vesicle content-mixing experiment using the quintuple mutant of Syt1 (Syt1_QM) that specifically disrupts the primary interface between Syt1 and the neuronal SNARE complex (Zhou et al., 2015). With reconstituted Syt1_QM, the stapled peptides had little effect on either Ca^2+^-independent or Ca^2+^-triggered fusion (Figs. 4E-J). The stapled peptides reduced vesicle association in the presence of Syt1_QM (Fig. 4D) which may be caused by binding of the stapled peptides to the polybasic region of Syt1 C2B, and thereby reducing Syt1/Syt2–membrane induced vesicle association in the absence of the primary interface. Together, these data suggest that the inhibitory effect of the stapled peptides on both Ca^2+^-independent and Ca^2+^-triggered fusion is due to specific interference with the primary interface between the SNARE complex and Syt1. Presumably, this interference occurs when the stapled peptides compete with binding of the SNARE complex to the C2B domain of Syt1 (Lai et al., 2022).

### Stapled peptides inhibit Ca^2+^-triggered vesicle fusion in the airway system

The residues in the primary interface are conserved across different species for synaptotagmins, SNAP-25, and Stx-1A homologues that are involved in fast synchronous release (Zhou et al., 2015). More generally, the similarity of residues involved in the primary interface between the neuronal system (neuronal SNAREs and Syt1 C2B) and the airway system (airway SNAREs VAMP8, Stx3, SNAP-23, and Syt2 C2B) is also high (Lai et al., 2022). Specifically, the residues that are at or near the primary interface are identical for Syt1/Syt2 except for V292C and identical for SNAP-25A/SNAP-23 except for K40Q, L47I, and V48T.

To confirm if these stapled peptides have a similar inhibitory activity for the airway system, we tested the peptides with a single-vesicle content-mixing assay with reconstituted airway SNAREs and Syt2 (Fig. 5A-B). Two types of vesicles were reconstituted: plasma-membrane mimicking vesicles with reconstituted Stx3 and SNAP-23 (referred to as airway PM vesicles) and vesicles with reconstituted VAMP8 and Syt2 that mimic mucin-containing secretory granules (referred to as SG vesicles). As expected, and consistent with the results with the neuronal system, the peptides showed no effect on vesicle association (Fig. 5C). In contrast, the stapled peptides strongly reduced Ca^2+^-triggered content mixing (Fig. 5F-I), along with mild inhibition of Ca^2+^-independent fusion (Fig. 5D-E). As a control, the unstapled P0 peptide showed no effect on either Ca^2+^-independent or Ca^2+^-triggered fusion. Similar to the neuronal case, when Syt2 was omitted, the stapled peptides showed no significant effect (Fig. 5J-L), again corroborating the notion that the stapled peptides function by interfering with the primary interface. Together, these results suggest that the inhibition activity of the stapled peptides is conserved for both the neuronal and airway systems.

**Figure 5.**
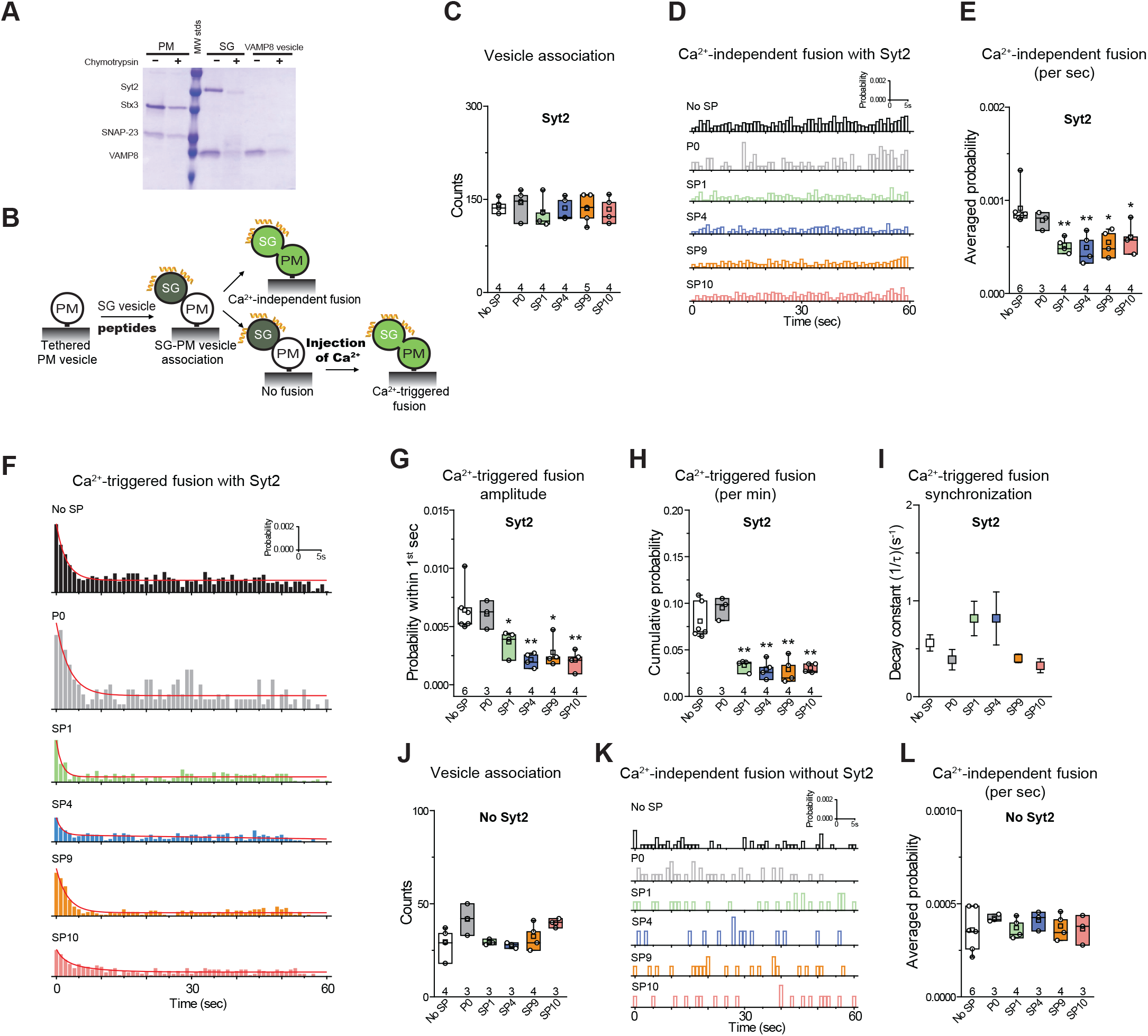
Stapled peptides inhibit Ca^2+^-triggered content mixing with reconstituted airway epithelial SNAREs and Syt2. **A,** SDS-PAGE of airway PM and SG vesicles with reconstituted airway SNAREs and Syt2. For details of the reconstitution and lipid compositions, see Methods in (Lai et al., 2022). **B,** Schematic of the single vesicle content mixing assay. Airway PM: plasma membrane mimic vesicles with reconstituted airway Stx3 and SNAP-23; SG: secretory granule mimics with reconstituted VAMP8 and Syt2. After SG - airway PM vesicle association, vesicle pairs either undergo Ca^2+^-independent fusion or remain associated until fusion is triggered by Ca^2+^ addition. 10 μM of each specific stapled peptide was added together with SG vesicles and was present during all subsequent stages. **C,** Effects of stapled peptides on vesicle association. **D,** Corresponding Ca^2+^-independent fusion probabilities. **E,** Corresponding average probabilities of Ca^2+^-independent fusion events per second (left to right: ** p = 0.0062, ** p = 0.0082, * p = 0.014, * p = 0.034). **F,** Corresponding Ca^2+^-triggered fusion probabilities. (**G-I**) Corresponding Ca^2+^-triggered fusion amplitudes of the first 1-sec time bin upon 500 μM Ca^2+^-injection (**G**) (left to right: * p = 0.036, ** p = 0.0033, * p = 0.012, ** p = 0.0037), the cumulative Ca^2+^-triggered fusion probability within 1 min. (**H**) (left to right: ** p = 0.0017, ** p = 0.0014, ** 0.0018, ** p = 0.0011), and the decay rate (1/τ) of the Ca^2+^-triggered fusion histogram (**I**). **J-L,** the stapled peptides have no effect on vesicle fusion mediated by airway SNAREs alone. **J,** Effects of 10 μM of each of the specified peptides on vesicle association using the assay described above. **K,** Corresponding Ca^2+^-independent fusion probabilities. **L,** Corresponding average probabilities of Ca^2+^-independent fusion events per second. Panels **C**, **E**, **G**, **H, J, L** show box plots and data points for n (indicated below each box plot) independent repeat experiments (Supplementary Table S1): the whiskers show the min and max values (excluding outliers), the box limits are the 25th and 75th percentiles, the square point denotes the mean value, and the center line denotes the median. Decay constants (boxes) and error estimates (bars) in panel **I** computed from the covariance matrix upon fitting the corresponding histograms combining all repeats with a single exponential decay function using the Levenberg-Marquardt algorithm. * p < 0.05, ** p < 0.01 by Student’s t-test, compared to the experiment without the stapled peptides.

## Discussion

We designed a series of hydrocarbon-stapled peptides that interfere with the primary SNARE/synaptotagmin interface and consequently disrupt Ca^2+^-triggered exocytosis (Fig. 1). We found that hydrocarbon staples increase the α-helical conformation of the peptide. The stapled peptides with the hydrocarbon-staple to bridge three amino acids (substitution positions *i* and *i*+4) exhibited better helical stabilization with the helical content up to 86% than the other designs (Fig. 1D,E). The stapled peptides (SP1, SP4, SP9, SP10) that consistently exhibit large inhibition in lipid mixing assay also have the largest α-helical content (Figs. 1,2). We therefore focused on these four stapled peptides in subsequent studies using a more advanced single-vesicle content-mixing assay that is capable of distinguishing between vesicle association, lipid-mixing, and content-mixing (Diao et al., 2012; Kyoung et al., 2011; Lai et al., 2022). The four stapled peptides reduced both Ca^2+^-triggered and Ca^2+^-independent fusion with reconstituted neuronal SNAREs and Syt1 (Fig. 3). When Syt1 was omitted, or replaced by a Syt1 mutant (Syt1_QM) that disrupts the primary interface (Zhou et al., 2015), there was no significant effect of the these stapled peptides on single vesicle fusion (Fig. 4), suggesting that the inhibitory activity of these peptides is related to the primary interface. The observed inhibition of Ca^2+^-triggered fusion is likely caused by stapled peptide binding to Syt1 in competition with the primary interface (Lai et al., 2022).

Next, we tested the selected stapled peptides with a reconstituted single vesicle fusion assay using airway SNAREs (VAMP8, Stx3, SNAP-23) and Syt2. The four stapled peptides also specifically inhibited Ca^2+^-triggered content mixing while the P0 control peptide showed no effect (Fig. 5). Interestingly, the effect of the stapled peptides on Ca^2+^-independent fusion was more modest compared to the neuronal case (compare Figs. 4, 5). For a more complete reconstitution involving Munc18 and Munc13 and using one of the stapled peptides (SP9), the effect on Ca^2+^-triggered fusion was also more pronounced than the effect on Ca^2+^-independent fusion (Lai et al., 2022). A possible explanation is that the airway system might lack complexin, a factor that both inhibits Ca^2+^-independent fusion, and activates Ca^2+^-triggered fusion (Lai et al., 2014; Maximov et al., 2009). Thus, since our “simple” reconstitution with reconstituted SNAREs and Syt1 (Figs. 2–4) lacks complexin, we may observe an enhanced effect of the stapled peptides on Ca^2+^-independent fusion, whereas for the airway system this function of complexin may not be required. In this context, it will also be desirable to test other sequences and strategies in future work, including the SNAP-23 sequence itself that might strengthen (or weaken) the interaction with Syt1 or Syt2 considering the K40Q, L47I, and V48T substitutions. We also note that Ca^2+^-independent fusion as observed with our assays may be distinct from spontaneous neurotransmitter release and baseline mucin secretion since they may use other high-affinity Ca^2+^-sensors (Kavalali, 2015).

In summary, our results indicate that the primary interface between the airway SNARE complex and Syt2 is also essential for Ca^2+^-triggered exocytosis. Moreover, the biological function of the primary interface between SNAREs and synaptotagmins is conserved for both synaptic neurotransmitter release and airway mucin secretion. We recently tested one of the stapled peptides (SP9) in cultured human airway epithelial cells and mouse airway epithelium by conjugation to cell penetrating peptides where it markedly and specifically reduced stimulated mucin secretion in both systems, and substantially attenuated mucus occlusion of mouse airways (Lai, Nature, 2022). Taken together, stapled peptides that disrupt Ca^2+^-triggered membrane fusion may allow therapeutic modulation of mucin secretory pathways and neurotransmitter release. Moreover, our inhibition strategy could be useful for a wide range of Ca^2+^-triggered endocrine and exocrine secretion systems.

## Acknowledgements

We thank Roberto Adachi, Melissa Singh, Thomas Südhof, Tony Velkov, Chuchu Wang for stimulating discussions, Heawon Chong and colleagues at Vivitide Inc. for their expertise in peptide synthesis and collaborative efforts, the National Institutes of Health (NIH) (R37MH63105 to A.T.B.; R01 HL129795 and R21 AI137319 to B.F.D.), and the Cystic Fibrosis Foundation (DICKEY18G0 and DICKEY19P0) for support. The contents of this publication are solely the responsibility of the authors and do not necessarily represent the official views of NIGMS or NIH. This article is subject to HHMI’s Open Access to Publications policy. HHMI lab heads have previously granted a nonexclusive CC BY 4.0 license to the public and a sublicensable license to HHMI in their research articles. Pursuant to those licenses, the author-accepted manuscript of this article can be made freely available under a CC BY 4.0 license immediately upon publication.

## Author Contributions

Y.L., B.F.D., and A.T.B. designed experiments. Y.L. performed fusion and CD experiments. Y.L., P.J., B.C., and Q.Z. performed inhibitor design. J.L. and J.P. wrote Matlab scripts to analyse the fusion experiments. R.A.P. and L.E. assisted with protein purification. Y.L., B.F.D., and A.T.B. wrote the manuscript.

## Methods

### Protein expression and purification

We used the same constructs and protocols to purify cysteine-free syntaxin-1A (Stx1), syntaxin-3 (Stx3), SNAP-25A, SNAP-23, VAMP2, VAMP8, Syt-1, Syt-2, and Syt-2_QM, as described in refs. (Lai et al., 2022, 2017). The protein sample concentrations were measured by UV absorption at 280 nm, aliquots were flash frozen in liquid nitrogen and stored at −80 °C.

### Peptide synthesis

All the stapled peptides (SP1, SP2, SP3, SP4, SP5 SP6, SP7, SP8, SP9, SP10, SP11, SP12) and the non-stapled peptide P0 were synthesized and purified by Vivitide (formerly New England Peptide, Gardner, USA). Peptide synthesis was carried out using solid phase peptide synthesis and Fmoc chemistry similar to ref. (Lai et al., 2022). Briefly, these peptides were cleaved using trifluoroacetic acid and standard scavengers. The peptides were purified using reverse phase high pressure liquid chromatography. For the stapled peptides, α,α-disubstituted non-natural amino acids of olefinic side chains were synthesized (*S*5 – *S* stereochemistry, bridging 5 amino acids; *R8* – *R* stereochemistry, bridging 8 amino acids). The hydrocarbon-staples were made via Grubbs catalyst (Schafmeister et al., 2000). For all stapled peptides, the N-termini were acetylated and the C-termini amidated. For more details of the synthesis see ref. (Lai et al., 2022).

All peptides were purified to > 90-95% purity, and quality control was performed by liquid chromatography mass spectrometry (LC-MS) by the manufacturer (Supplementary Figs. S1-S12). Subsequently, the peptides were lyophilized and shipped. The LC-MS quality control data indicate that the peptides have the predicted molecular weight according to their chemical composition, suggesting that the synthesized peptides have the correct sequence and properly formed staples.

For each group of experiments, aliquots of peptide powder were directly dissolved in the specified buffers at ~ 1 mM concentration using a vortexer, and then diluted to the specified peptide concentrations. The concentration of the stock solution was confirmed by absorption measurement at 205 nm using a Nanodrop instrument (Thermo Fisher, Inc.).

### CD spectroscopy

The CD spectra were measured with an AVIV stop-flow CD spectropolarimeter at 190 to 250 nm using a cell with a 1 mm path-length. The sample containing 100 mM of synthesized peptides in PBS buffer (137 mM NaCl, 2.7 mM KCl, 10 mM Na2HPO4, and 2 mM KH2PO4, pH 7.4) was measured at 20 °C. For the correction of the baseline error, the signal from a blank run with PBS buffer was subtracted from all the experimental spectra. The α-helical content of each peptide was calculated by dividing the mean residue ellipticity [φ]222obs by the reported [φ]222obs for a model helical decapeptide (Yang et al., 1986).

### Vesicle reconstitution

For the ensemble lipid-mixing assay, the lipid composition of SV vesicles was phosphatidylcholine (PC) (46%), phosphatidylethanolamine (PE) (20%), phosphatidylserine (PS) (12%), cholesterol (20%), and 1,1’-dioctadecyl-3,3,3’,3’-tetramethylindodicarbocyanine perchlorate (DiD) (2%). For PM vesicles the lipid composition was Brain Total Lipid Extract (Avanti Polar Lipids) supplemented with 3.5 mol% PIP2, 0.1 mol% biotinylated phosphatidylethanolamine (PE) and 1,1’-dioctadecyl-3,3,3’,3’-tetramethylindocarbocyanine perchlorate (DiI).

For single-vesicle content-mixing assays, the lipid composition of SV vesicles, SG vesicles, VAMP2-only, or VAMP8-only vesicles was phosphatidylcholine (PC) (48%), phosphatidylethanolamine (PE) (20%), phosphatidylserine (PS) (12%), and cholesterol (20%). For PM vesicles, the lipid composition was Brain Total Lipid Extract (Avanti Polar Lipids) supplemented with 3.5 mol% PIP2, and 0.1 mol% biotinylated phosphatidylethanolamine (PE). The detailed reconstitution protocols for the vesicles are described in refs. (Lai et al., 2022, 2017).

### Ensemble lipid mixing

Protein-reconstituted PM and SV vesicles were mixed at a molar ratio of 1:1, both at 0.1 mM lipid concentration. In the ensemble lipid mixing assay, the SNARE protein concentrations are ~ 0.5 μM, while the peptide concentrations are 10 μM and 100 μM, so the peptide:SNARE ratios are 1:20 or 1:200, respectively. Lipid-mixing is measured as the fluorescent emission (at 670 nm) of DiD dyes in SV vesicles via fluorescence resonance energy transfer (FRET) from DiI dyes in PM vesicles that were excited by 530 nm laser light. The fluorescence intensities were monitored in two channels at 570 nm and 670 nm, respectively. Fluorescence changes were recorded with a Varian Cary Eclipse model fluorescence spectrophotometer using a quartz cell of 100 μl with a 5 mm path length. All measurements were performed at 35°C.

### Single vesicle-vesicle content mixing assay, instrument setup, and data analysis

Details for the single-vesicle content-mixing assay are described in (Lai et al., 2022). In the single vesicle content mixing assay, the peptide concentration is 10 μM in the chamber. The PM vesicles are tethered to the surface with ~ 100 syntaxin molecules per vesicle (Kyoung et al., 2011), so considering the two-dimensional local concentration effect, the peptide:SNARE ratio is expected to be very high.

## Quantification and Statistical Analysis

Origin, Matlab, and Prism were used for the generation of all curves and graphs. The single-vesicle content-mixing experiments were repeated at least three times with different protein preps and vesicle reconstitutions (Supplementary Table S1) and properties were calculated as mean ± SEM. The two-tailed Student’s t-test was used to test statistical significance in Figs. 3, 4, 5 with respect to the specified reference experiment.

## Software and code

The data for the ensemble lipid mixing experiments were collected by a program provided by the manufacturer of the Cary Eclipse instrument. The data for the single-vesicle content-mixing experiments were collected by using the smCamera program developed by Taekjip Ha, Johns

Hopkins University, Baltimore. For the ensemble lipid mixing experiments, the single-vesicle content-mixing experiments and the circular dichroism experiments, OriginPro 8 and Matlab-2021b were used. Pymol 2.5.1 was used for visualization. MATLAB analysis scripts for the single-vesicle fusion experiments and for the smCamera file conversions are available in the Zenodo repository https://doi.org/10.5281/zenodo.6370585.

## Data availability

The datasets presented in this study can be found in the Dryad repository https://doi.org/10.5061/dryad.x3ffbg7mp.

## Supplementary material

**Supplementary Table S1. Data summary table for the single vesicle fusion experiments**

**Supplementary Figure S1.** Quality control data for the P0 peptide provided by Vivitide Inc.

**Supplementary Figure S2.** Quality control data for the P1 peptide provided by Vivitide Inc.

**Supplementary Figure S3.** Quality control data for the P2 peptide provided by Vivitide Inc.

**Supplementary Figure S4.** Quality control data for the P3 peptide provided by Vivitide Inc.

**Supplementary Figure S5.** Quality control data for the P4 peptide provided by Vivitide Inc.

**Supplementary Figure S5.** Quality control data for the P5 peptide provided by Vivitide Inc.

**Supplementary Figure S6.** Quality control data for the P6 peptide provided by Vivitide Inc.

**Supplementary Figure S7.** Quality control data for the P7 peptide provided by Vivitide Inc.

**Supplementary Figure S8.** Quality control data for the P8 peptide provided by Vivitide Inc.

**Supplementary Figure S9.** Quality control data for the P9 peptide provided by Vivitide Inc.

**Supplementary Figure S10.** Quality control data for the P10 peptide provided by Vivitide Inc.

**Supplementary Figure S11.** Quality control data for the P11 peptide provided by Vivitide Inc.

**Supplementary Figure S12.** Quality control data for the P12 peptide provided by Vivitide Inc.

