## Supplementary Figure S8 for "Screening of hydrocarbon-stapled peptides for inhibition of calcium-triggered exocytosis"

### Certificate of Analysis

| **Sequence:** [Cyc(4,11)]Ac-SKD(R8)GIRTLV(S5)LDEQGEQL-amide | | |
|---|---|---|
| **Peptide Name:** | | **Date:** 8/1/2017 |
| **Order#:** P611359 | **Lot#:** LB1364 | **Amount:** 5.1mg |

**Quality Control Specifications:**

| QC Test | QC Specifications | Results |
|---|---|---|
| Purity by HPLC | ≥**90%** by percent area | **Pass** |
| Mass Identification by Mass Spectral Analysis | Calculated Mass within 0.1% of Molecular Weight: **2235** | **Pass** |
| Concentration/ Net Peptide | Amino Acid Analysis (AAA) determining original concentration/net peptide content. | **N/A** |

**Product:** Research Grade Custom Peptide containing traces of Trifluoroacetate (TFA) salts.

**Formulation:**
Final concentration: N/A
Final form: Dry

**Stability and Conditions:** Refer to the Quality Control Detail Information on our website at www.newenglandpeptide.com/support/quality-control-information . As always, NEP has individual batch records stored electronically for each peptide that includes traceable lot numbers of raw materials used during synthesis. Should you require this information, with your peptide lot number.

**Notes (if applicable):**

AD
**Approval/Initials**

### Peptide QC Report LB1364 12-15

| | |
|---|---|
| Analysis Name | D:\Data\LB1364 12-15_141922_P1-A-4_01_71062.D |
| Sample Name | LB1364 12-15 |
| Method | APRIL20171.2mLperMIN_NEPO<br>AHIGH_71062.m |
| Instrument | amaZon SL |

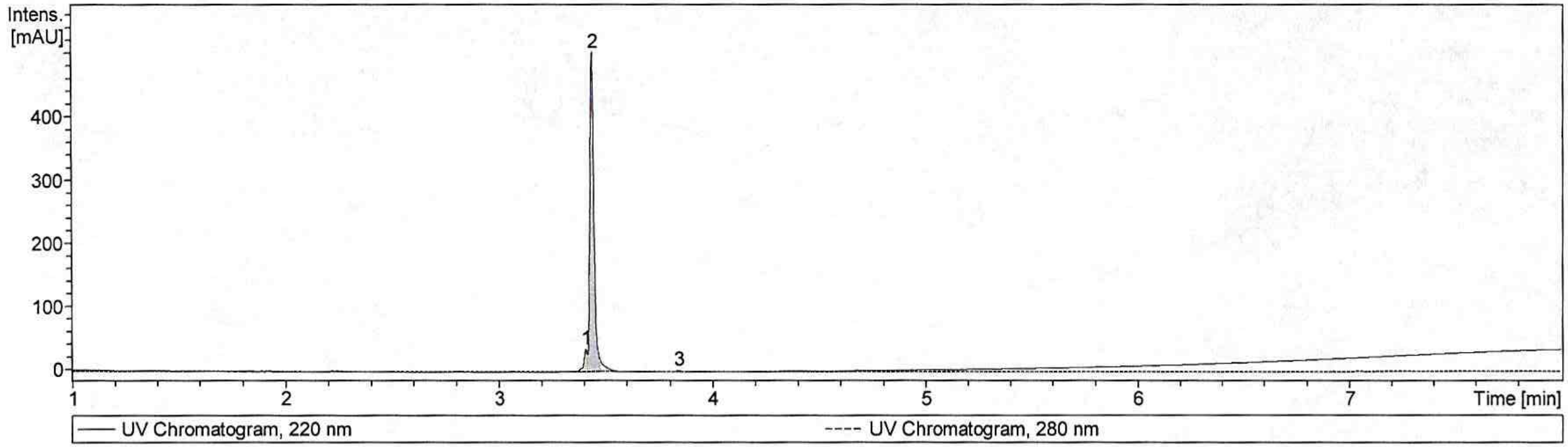


| Target Mass | Meas. Mass | Expec. Mass | Delt. Mr [Da] | Intensity | Area | Area Fraction [%] |
|---|---|---|---|---|---|---|
| Cmpd 2; 3.44 min; Pep Mr: 2234.25 | 2234.25 | 2235.00 | -0.75 | 498 | 632 | 93.3 |

| # | RT [min] | Area | Area Frac. % |
|---|---|---|---|
| 1 | 3.41 | 41.8613 | 6.18 |
| 2 | 3.44 | 632.2511 | 93.27 |
| 3 | 3.84 | 3.7519 | 0.55 |

Cmpd 2; 3.44 min; Pep Mr: 2234.25

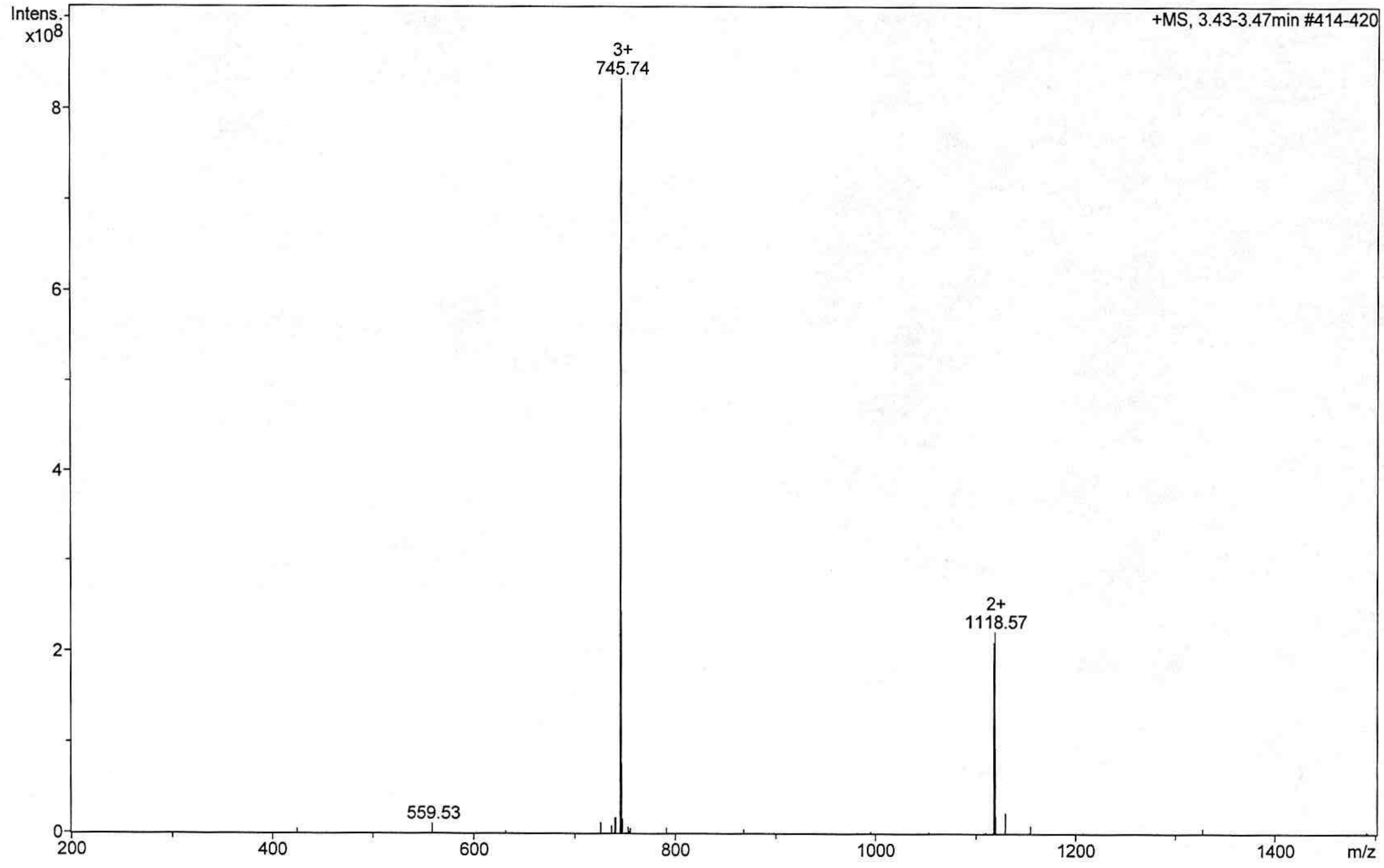
