## Supplementary Figure S9 for "Screening of hydrocarbon-stapled peptides for inhibition of calcium-triggered exocytosis"

new england peptide™

### Certificate of Analysis

| **Sequence:** [Cyc(3,7;10,14)]Ac-EE(S5)KDA(S5)IR(S5)LVM(S5)DEQC-amide | | |
|---|---|---|
| **Peptide Name:** | | **Date:** 8/7/2017 |
| **Order#:** P611359 | **Lot#:** LB1544 | **Amount:** 5.4mg |

**Quality Control Specifications:**

| QC Test | QC Specifications | Results |
|---|---|---|
| Purity by HPLC | ≥**90%** by percent area | **Pass** |
| Mass Identification by Mass Spectral Analysis | Calculated Mass within 0.1% of Molecular Weight: **2222** | **Pass** |
| Concentration/ Net Peptide | Amino Acid Analysis (AAA) determining original concentration/net peptide content. | **N/A** |

**Product:** Research Grade Custom Peptide containing traces of Trifluoroacetate (TFA) salts.

**Formulation:**
Final concentration: N/A
Final form: Dry

**Stability and Conditions:** Refer to the Quality Control Detail Information on our website at www.newenglandpeptide.com/support/quality-control-information . As always, NEP has individual batch records stored electronically for each peptide that includes traceable lot numbers of raw materials used during synthesis. Should you require this information, with your peptide lot number.

**Notes (if applicable):**

**Approval/Initials**

*For Science... From Science.*

New England Peptide Inc., 65 Zub Lane, Gardner, MA 01440 ▪ **Phone** 888-343-5974 ▪ **Fax** 978-630-0021

**www.NewEnglandPeptide.com**

### Peptide QC Report LB1544 25-43

| | |
|---|---|
| Analysis Name | D:\Data\LB1544 25-43_143070_P1-E-5_01_71582.D |
| Sample Name | LB1544 25-43 |
| Method | APRIL20171.2mLperMIN_NEPO AHIGH_71582.m |
| Instrument | amaZon SL |

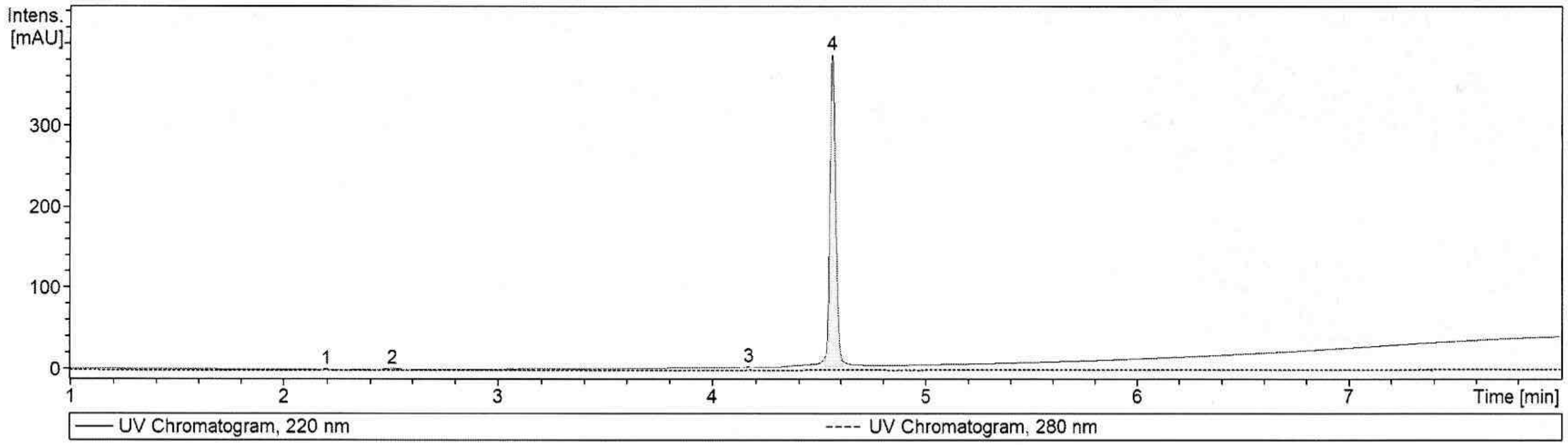


| Target Mass | Meas. Mass | Expec. Mass | Delt. Mr [Da] | Intensity | Area | Area Fraction [%] |
|---|---|---|---|---|---|---|
| Cmpd 4; 4.56 min; Pep Mr: 2221.04 | 2221.04 | 2222.00 | -0.96 | 386 | 713 | 98.8 |

| # | RT [min] | Area | Area Frac. % |
|---|---|---|---|
| 1 | 2.20 | 1.7029 | 0.24 |
| 2 | 2.50 | 5.3905 | 0.75 |
| 3 | 4.17 | 1.2394 | 0.17 |
| 4 | 4.56 | 712.5846 | 98.84 |

### Peptide QC Report LB1544 25-43

**Cmpd 4; 4.56 min; Pep Mr: 2221.04**

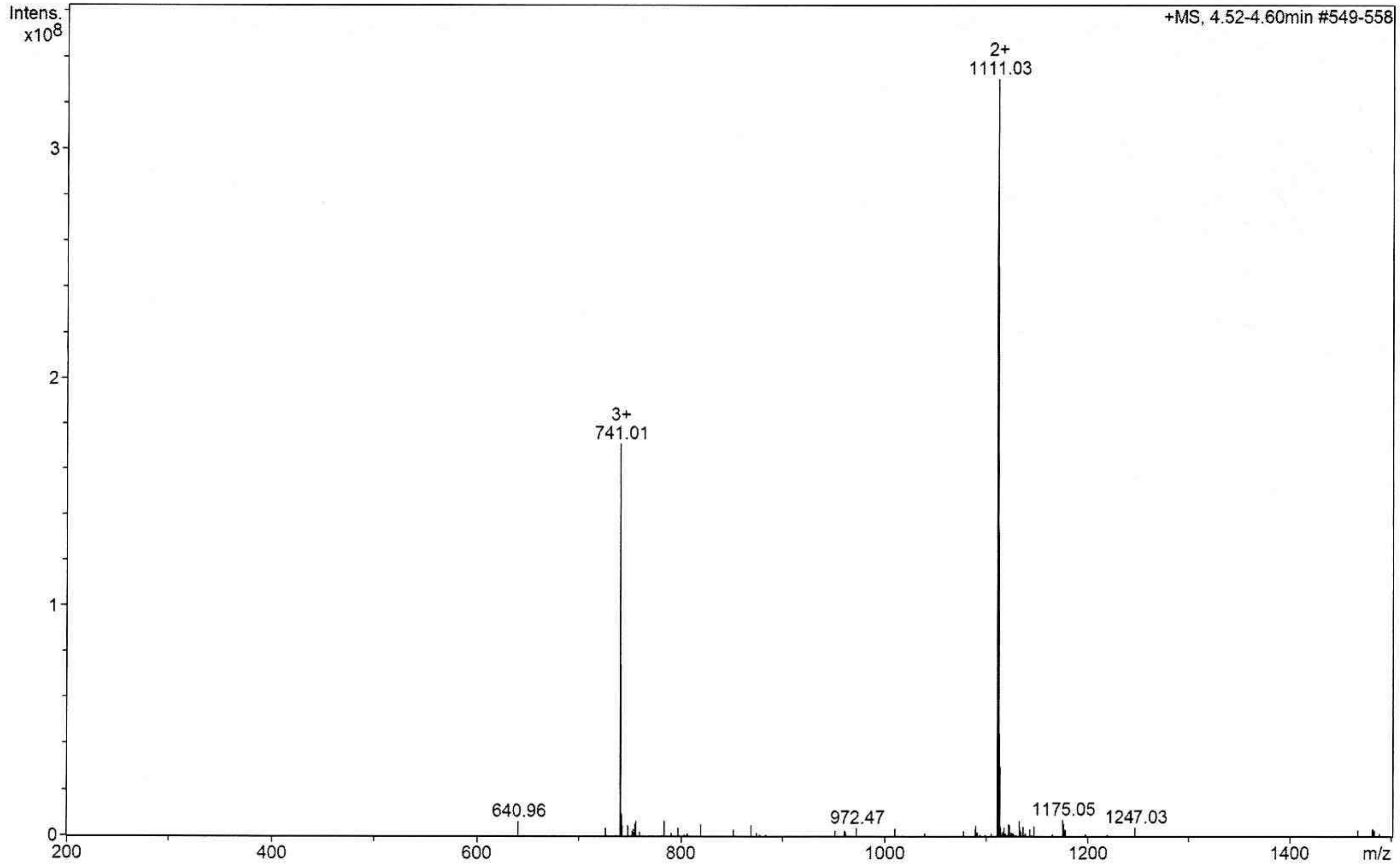
