## Supplementary Figure S10 for "Screening of hydrocarbon-stapled peptides for inhibition of calcium-triggered exocytosis"

nep new england peptide™

### Certificate of Analysis

| **Sequence:** [Cyc(4,8;11,15)]Ac-SKD(S5)GIR(S5)LV(S5)LDE(S5)GEQL-amide | | |
|---|---|---|
| **Peptide Name:** | | **Date:** 8/7/2017 |
| **Order#:** P611359 | **Lot#:** LB1542 | **Amount:** 5.1mg |

**Quality Control Specifications:**

| QC Test | QC Specifications | Results |
|---|---|---|
| Purity by HPLC | ≥90% by percent area | **Pass** |
| Mass Identification by Mass Spectral Analysis | Calculated Mass within 0.1% of Molecular Weight: **2215** | **Pass** |
| Concentration/ Net Peptide | Amino Acid Analysis (AAA) determining original concentration/net peptide content. | **N/A** |

**Product:** Research Grade Custom Peptide containing traces of Trifluoroacetate (TFA) salts.

**Formulation:**
Final concentration: N/A
Final form: Dry

**Stability and Conditions:** Refer to the Quality Control Detail Information on our website at www.newenglandpeptide.com/support/quality-control-information . As always, NEP has individual batch records stored electronically for each peptide that includes traceable lot numbers of raw materials used during synthesis. Should you require this information, with your peptide lot number.

**Notes (if applicable):**

**Approval/Initials**



New England Peptide Inc., 65 Zub Lane, Gardner, MA 01440 ■ **Phone** 888-343-5974 ■ **Fax** 978-630-0021

**www.NewEnglandPeptide.com**

### Peptide QC Report LB1542 5-24

| | |
|---|---|
| Analysis Name | D:\Data\LB1542 5-24_143061_P1-D-8_01_71578.D |
| Sample Name | LB1542 5-24 |
| Method | APRIL20171.2mLperMIN_NEPO AHIGH_71578.m |
| Instrument | amaZon SL |

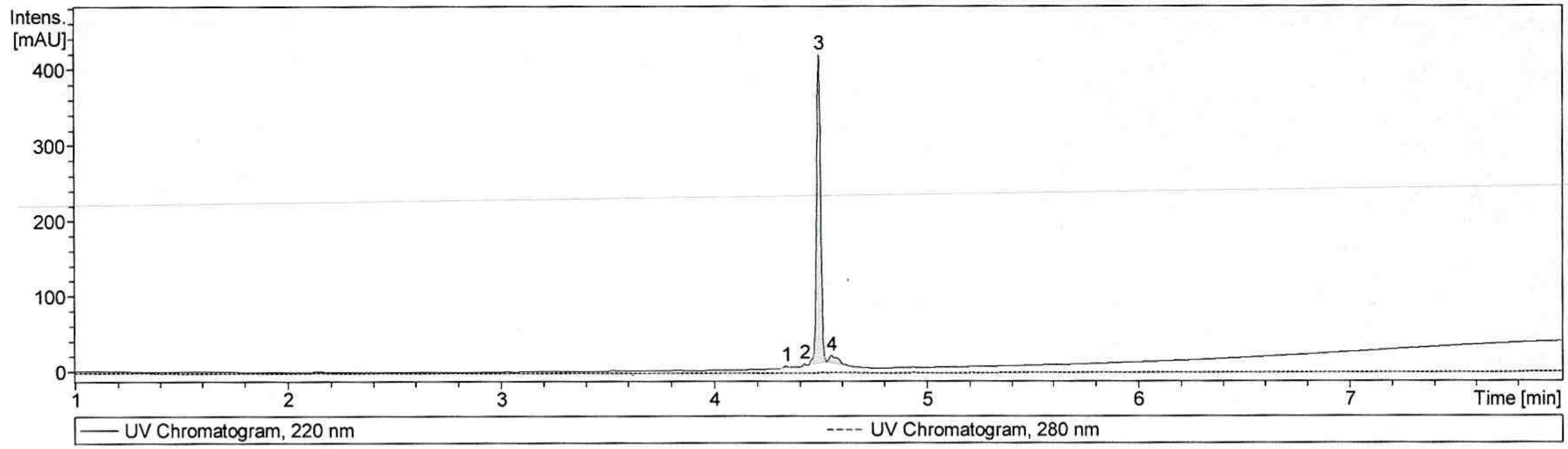


| Target Mass | Meas. Mass | Expec. Mass | Delt. Mr [Da] | Intensity | Area | Area Fraction [%] |
|---|---|---|---|---|---|---|
| Cmpd 3; 4.49 min; Pep Mr: 2214.15 | 2214.15 | 2215.00 | -0.85 | 418 | 545 | 94.3 |

| # | RT [min] | Area | Area Frac. % |
|---|---|---|---|
| 1 | 4.33 | 2.7407 | 0.47 |
| 2 | 4.42 | 2.8883 | 0.50 |
| 3 | 4.49 | 544.5035 | 94.29 |
| 4 | 4.55 | 27.3173 | 4.73 |

### Peptide QC Report LB1542 5-24

Cmpd 3; 4.49 min; Pep Mr: 2214.15

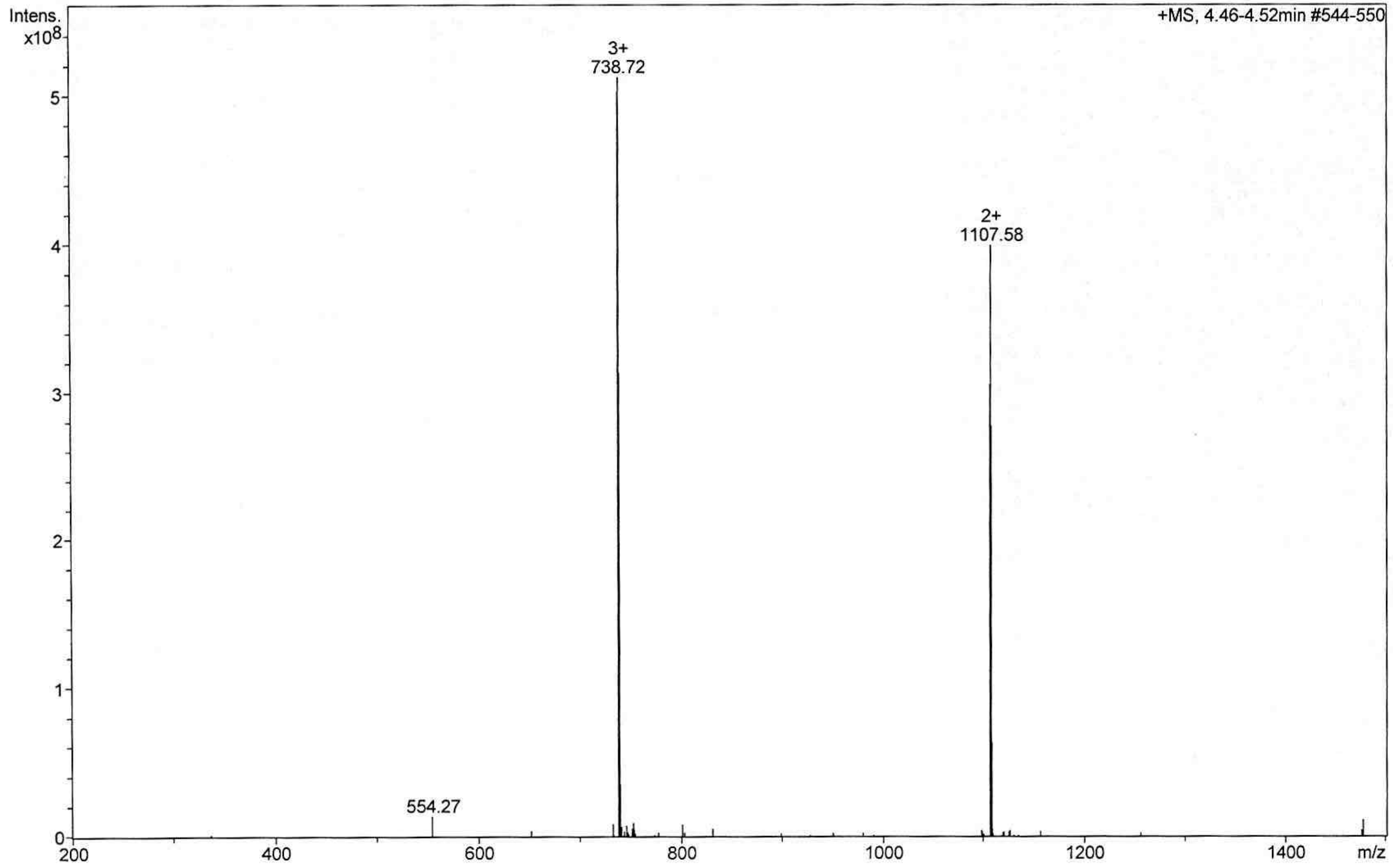
