## Supplementary Table S1 for "Screening of hydrocarbon-stapled peptides for inhibition of calcium-triggered exocytosis"

**Supplementary Table 1. Data summary table for the single vesicle fusion experiments**

| Tested peptides or conditions | Ca^2+^-independent  fusion events | Ca^2+^-triggered  fusion events | Number of analyzed  vesicle pairs | Repeats |
| --- | --- | --- | --- | --- |
| neuronal SNAREs and Syt1 (related to Fig. 3) | | | | |
| None | 750 | 846 | 9557 | 5 |
| P0 | 493 | 591 | 6543 | 3 |
| SP1 | 235 | 210 | 7063 | 4 |
| SP4 | 261 | 210 | 8773 | 4 |
| SP9 | 165 | 272 | 7122 | 4 |
| SP10 | 263 | 495 | 8616 | 4 |
| neuronal SNAREs only (related to Fig. 4) | | | | |
| None | 410 |  | 6980 | 8 |
| P0 | 104 |  | 2020 | 4 |
| SP1 | 245 |  | 5650 | 4 |
| SP4 | 356 |  | 6246 | 4 |
| SP9 | 336 |  | 6546 | 4 |
| SP10 | 341 |  | 6198 | 4 |
| neuronal SNAREs and Syt1_QM (related to Fig. 4) | | | | |
| None | 320 | 384 | 6480 | 3 |
| SP1 | 226 | 282 | 5044 | 3 |
| SP4 | 222 | 248 | 4878 | 3 |
| SP9 | 242 | 260 | 2989 | 3 |
| SP10 | 252 | 367 | 5329 | 3 |
| airway SNAREs and Syt2 (related to Fig. 5) | | | | |
| None | 461 | 657 | 8377 | 6 |
| P0 | 113 | 219 | 2306 | 3 |
| SP1 | 251 | 275 | 8126 | 4 |
| SP4 | 237 | 231 | 8017 | 4 |
| SP9 | 274 | 250 | 8445 | 4 |
| SP10 | 327 | 273 | 8925 | 4 |
| airway SNAREs only (related to Fig. 5) | | | | |
| None | 54 |  | 2578 | 6 |
| P0 | 38 |  | 1448 | 3 |
| SP1 | 26 |  | 1094 | 4 |
| SP4 | 16 |  | 660 | 3 |
| SP9 | 22 |  | 966 | 4 |
| SP10 | 18 |  | 840 | 3 |

Among each repeat experiment there are at least three different protein preps and vesicle reconstitutions, so the variations observed in the bar charts reflect sample variations as well as variations among different flow chambers. For the definition of the repeat experiments see (Lai et al., 2022)
